# Protein Language Model for Prediction of Subcellular Localization of Protein Sequences from Gram-negative bacteria (ProtLM.SCL)

**DOI:** 10.1101/2022.12.16.520742

**Authors:** Gurpreet Singh, Ravi Tyagi, Anjana Singh, Shruti Kapil, Pratap Kumar Parida, Maria Scarcelli, Dan Dumitru, Nanda Kumar Sathiyamoorthy, Sanjay Phogat, Ahmed Essaghir

## Abstract

The prediction of bacterial protein Sub-Cellular Localization (SCL) is critical for antigen identification and reverse vaccinology, especially when determining protein localization in the lab is time consuming, expensive and not possible for all species. While PSORTb is one of the most widely used tool for predicting SCL, it has several limitations, including the tendency to label a large number of proteins as ‘Unknown’. To address these shortcomings, we present a protein language model capable of predicting the subcellular localization of a given protein (ProtLM.SCL) from gram-negative bacteria. By performing 10-fold cross validation on the PSORTb public data set, we demonstrate that ProtLM.SCL is more accurate and precise than PSORTb. When compared to empirically validated published data, our models also outperformed PSORTb, particularly when categorizing difficult occurrences.

## 1. Introduction

Bacterial infections are a leading cause of death and disease in humans. Antibiotics and vaccines are the first line of medical and prophylactic defence against bacterial infection, but antibiotic resistance is becoming more prevalent and rapidly spreading^1^. Numerous microbial diseases, including tuberculosis, pneumonia, septicemia, and childhood ear infections, are becoming antibiotic-resistant^2–4^. Bacteria’s primary attack/defence against their hosts are secretory and surface proteins. Certain surface proteins of bacteria can aid them in adhering to and invading host cells, as well as aid them in fighting off host defences, thereby contributing significantly to bacterial pathogenicity. Additionally, outer-membrane and surface-exposed proteins contribute to bacterial pathogenicity and interact with the host immune system, making them a good candidate for vaccines and drug targets^5,6^.

There are numerous wet lab methods available for validating the subcellular localization of a proteins, including fluorescence microscopy and mass spectrometry^7,8^. Some other methods such as fractionation by using gradient centrifugation or 2D gel electrophoresis are also commonly used for determining the subcellular localization of proteins^9^. However, these methods can be time consuming and costly, and may even require several iterations during the experimental validation process^10^.

The subcellular localization of a protein helps determine whether it is exposed to the surface and may hint towards its biological function^11^. A typical gram-negative bacterium is roughly composed of five subcellular compartments: cytoplasm, inner membrane, periplasm, outer membrane, and extracellular space^12^. It is critical that a protein is directed to the appropriate cellular compartment in order to accomplish its function. Thus, the availability of an advanced computational method capable of accurately and precisely predicting subcellular localization will be critical in identifying potential drug or vaccine targets. Further, predicting a protein’s subcellular localization becomes vital for bioinformaticians as the number of unannotated/unreviewed protein sequence data in UniProt increases^10^.

Recently, some *in-silico* methods have been developed to predict the subcellular localization of a protein^10^. These methods augment wet lab experimental approaches by enabling rapid prediction of surface exposition or identification of SCL of proteins, thereby reducing the time required for the development of novel drugs or vaccines^13^. These computational prediction methods are mostly based on amino acid composition, known target sequences/motifs, and sequence similarity^14^. One such tool that has been widely used is PSORTb^15,16^. Since 2003, it has been one of the most precise and accurate predictors for SCL available for single and batch sequence processing^16^. However, PSORTb is designed to be more precise than sensitive. As a result, several protein types are poorly predicted or annotated.

In this study, we present ProtLM.SCL, a natural language model capable of predicting the subcellular localization of a given amino acid sequence while achieving higher performance. This was accomplished by training a self-supervised language model for proteins that represents long-range and bidirectional context through self-attention and self-supervision, in which amino acid sequences were considered as letters that comprise the words and phrases describing a protein’s primary structure^17,18^. Thus, ProtLM.SCL predicts the surface exposure or subcellular localization of bacterial proteins using just primary sequences of proteins as an input.

Our main contribution in this field are three-folds. Firstly, we present a protein language model specifically trained on a large dataset of sequences from UniRef50 which may be also used to obtain sequence embeddings representing clusters of protein sequences^19^. Secondly, we propose a fine-tuned model, ProtLM.SCL, that exhibits state-of-the-art performance in predicting SCL over held-out clusters of experimentally validated sequences from publicly available data sets. Lastly, our findings indicate that, as a result of the enhanced performance of ProtLM.SCL, we overcame key coverage limitation for PSORTb, which classifies numerous proteins as ‘Unknown’.

## 2. Material & Methods

### Pretraining of general protein language model

We pretrained the general language model to learn amino-acid physio-chemical properties and co-occurrences in protein sequences using the UniRef50 dataset release version 2020_02 dated Apr 22, 2020 ^19^. It contained a total of 39,949,058 sequences that were stratified by organism to generate a dataset for language model training in the split ratio of 94.05% train, 4.95% valid, 1% test with no duplicates. The language modelling was accomplished using masked language modelling paradigm with cross-entropy as the loss function. We have used transformer based RoBERTa model styled architecture^20^. We used perplexity and cross-entropy loss to determine when to stop training the model and choose the most accurate model. The final values for the cross-entropy loss and perplexity were 2.36 and 10.59 respectively.

### Pre-processing of downstream dataset

We have utilized the gram-negative dataset, containing a total of 8,224 proteins, that was used to train PSORTb v.3.0^16^. We conducted two types of comparisons in this study, classifying a given protein sequence into (1) binary categories: Surface Exposed Proteins (SEP) *vs*. Non-SEP, and (2) multiple categories: cytoplasmic, cytoplasmic membrane, periplasmic, outer membrane, and extracellular.

For binary classification, we have treated outer membrane, outer membrane/extracellular, and extracellular classes as SEP and cytoplasmic, cytoplasmic/cytoplasmic membrane, and cytoplasmic membrane classes as Non-SEP. We excluded periplasmic proteins from this version of the model since they are ambiguous and needed additional curation to be classified into SEPs or Non-SEPs and got 7,726 proteins. The SEP and Non-SEP categories, respectively, contained 1,048 and 6,678 proteins.

For multi-class classification, the original dataset contained a total of nine classes i.e. cytoplasmic, cytoplasmic/cytoplasmic membrane, cytoplasmic membrane, cytoplasmic membrane/periplasmic, periplasmic, periplasmic/outer membrane, outer membrane, outer membrane/extracellular and extracellular. To build a dataset containing all distinct classes we assigned the proteins with two labels to both classes, hence creating a dataset with a total of five labels i.e. cytoplasmic, cytoplasmic membrane, periplasmic, outer membrane, and extracellular containing 5071, 1711, 498, 632, and 507 proteins respectively.

#### 2.1. Strategy for Data Splitting

We were mindful of the fact that proteins can contain identical patterns, which could result in data leakage if training-testing validation were performed randomly. As a result, we used CD-HIT clustering with a 50% similarity threshold to generate protein clusters. Clusters were assigned to 11 buckets in such a way that each cluster is uniquely associated with a single bucket. One of the buckets was used as a holdout set (11^th^ bucket), while the remaining 1-10 buckets were used for 10-fold cross validation training (**Figure 1**).

**Figure 1:**
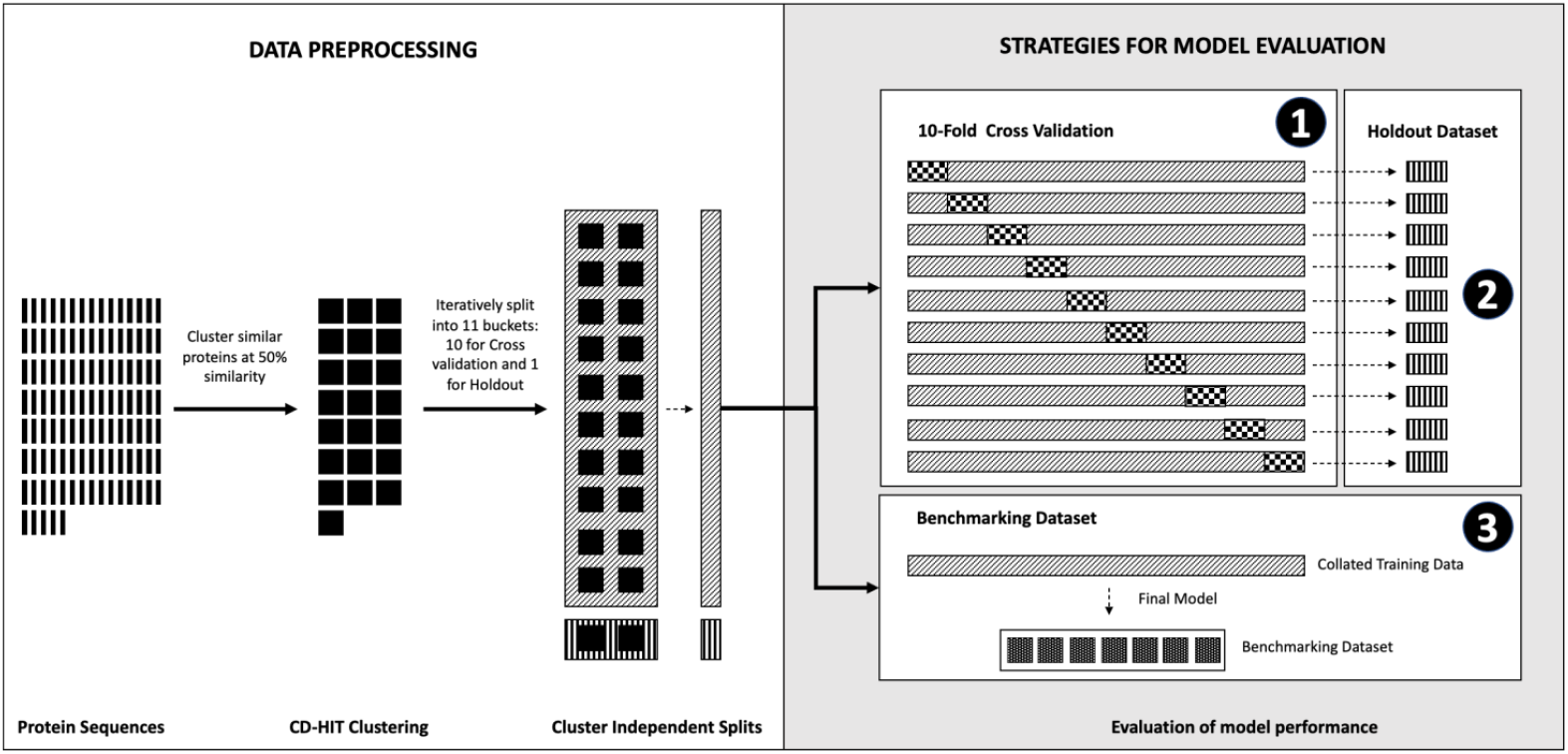
Overview of data pre-processing and strategies for model evaluation. We employed CD-HIT clustering with a 50% similarity criterion to construct protein clusters. Clusters were assigned to 11 buckets in such a way that each cluster is uniquely associated with a particular bucket. One of the buckets was utilized as a holdout set, while the remaining ten buckets were used for 10-fold cross validation training. We created each model for binary and multi-class classification using a one vs rest strategy and evaluated the performance of the models using three types of analysis: (1) **Cross-validation dataset:** We perform 10-fold cross-validation using protein sequences from 10 buckets and report these results as mean standard deviation across all folds (2) **Hold-out dataset:** We use protein sequences from all 10 buckets as the training set to create one ‘final model’ Each final model was then checked for overfitting using the 11th bucket as the hold-out dataset (3) **Benchmarking dataset:** We created a benchmarking dataset by collating protein sequences from seven experimentally validated datasets^16,21–26^. We report model performance on this dataset for each task using the ‘final model’ as an approximation for real-world use case

Label wise distribution of training data for binary & multi-class classification is given in **Table S1**. For binary classification, we obtained 11 buckets with an average of 702±30 proteins containing 12.54±2.54% of SEP proteins (**Table S2**). For multiclass classification, we obtained a total of 11 buckets with an average of 747±22 proteins containing 61.64±3.44% of cytoplasmic, 20.73±2.24% of cytoplasmic membrane, 6.00±1.26% of periplasmic, 7.64±1.36% of outer membrane and 6.18±1.08% of extracellular proteins (**Table S3**).

#### 2.2. Evaluation of model performance

For both binary and multi-class classification we created each model using one vs rest strategy and evaluated the performance of the models using three types of analysis

1. **Cross-validation dataset:** We performed 10-fold cross-validation by using protein sequences from 10 buckets and report these results as mean ± standard deviation across all folds.
2. **Hold-out dataset:** Here, we used protein sequences from all the 10 buckets as the training set to create one ‘final model’ each for binary and multi-class classification. Each final model was then checked for overfitting by using the 11^th^ bucket as the hold-out dataset.
3. **Benchmarking dataset:** We created a benchmarking dataset by collating protein sequences from seven experimentally validated datasets^16,21–26^ (**Table S4**). Detailed information regarding the benchmarking datasets is given in the supplementary file (S2). We report the model performance on this dataset for each task using the ‘final model’ as an approximation for real-world use cases. Here, we were mindful that some sequences could have been shared between this benchmarking dataset and the training dataset for language model. Hence, we pre-processed and removed the overlapping proteins. Finally, we had 3950 proteins for prediction of SEP *vs*. Non-SEP and 3810 proteins for prediction of SCL (**Table S4**). The standardized subcellular localization labels were added to the benchmark dataset according to the PSORTb nomenclature (**Table S5**). Details of distribution of SCL proteins localization wise on benchmark dataset is provided in **Figure 2**.

**Figure 2:**
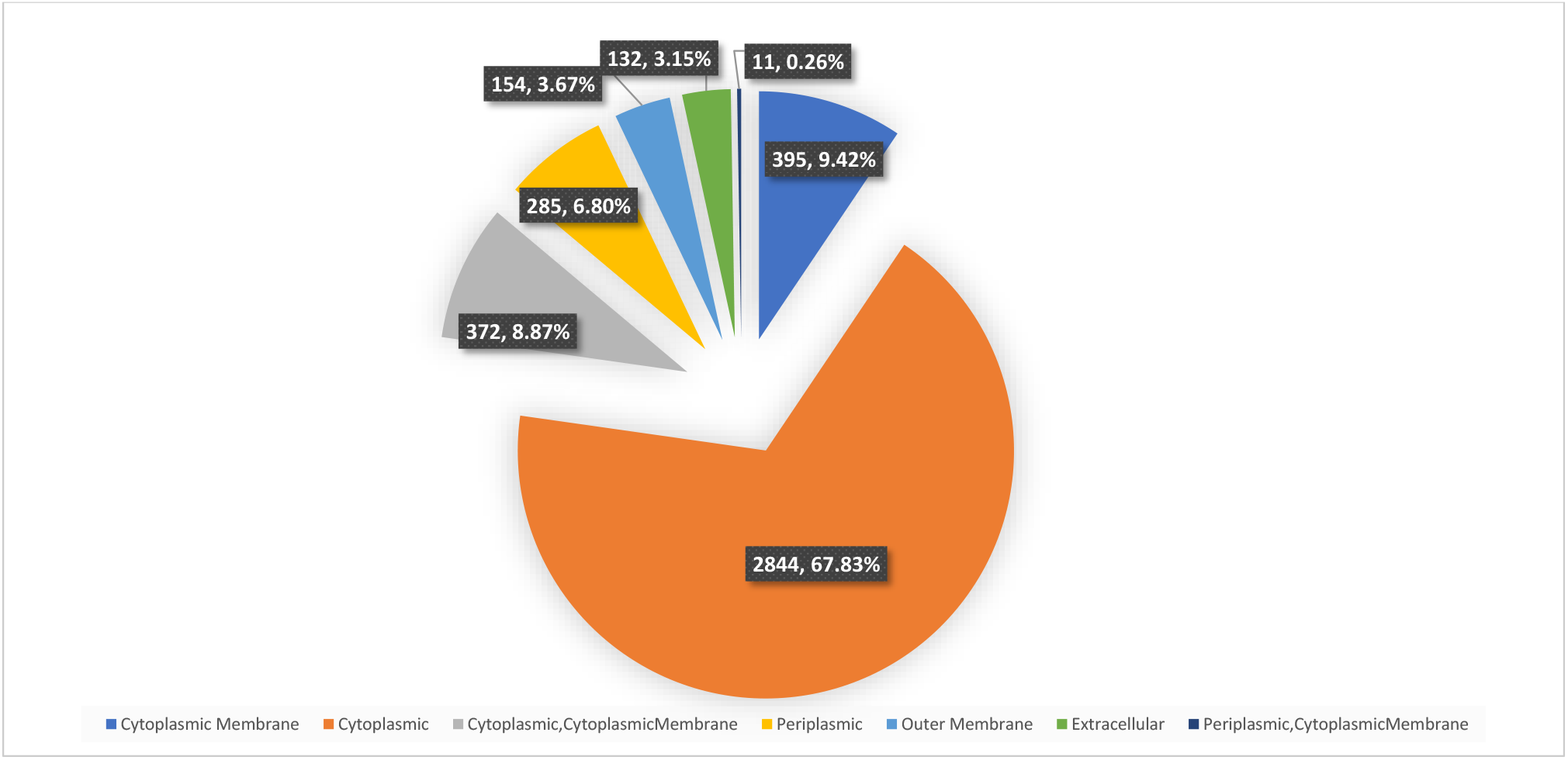
Class-wise distribution of the protein sequences for benchmarking dataset.

#### 2.3. Model Fine-tuning for downstream classification

The downstream model requires only primary protein sequences as an inputs, as illustrated in **Figure 3**. To perform binary classification, the pretrained language model was appended with a dense classifier layer and fine-tuned to specialize in classifying a given protein into SEP vs. Non-SEP as described in **section 2.2**. To accomplish multi-class classification, we trained five distinct instances of the language model, one for each of the five target labels using one vs. rest strategy, namely cytoplasmic, cytoplasmic membrane, periplasmic, outer membrane, and extracellular. It is also important to note that there were a total of 147 proteins which were assigned to multiple labels. To determine the final localization, we used the highest probability score derived from the output of the five models. Also, to check the improvement we calculated percentage gain (by MCC and Precision) of ProtLM.SCL over PSORTb using the formula; Highest-lowest/lowest*100. Additionally, if none of the models achieved a probability of >0.5, the prediction for such protein an input is considered an error and labelled as ‘Unknown’.

**Figure 3:**
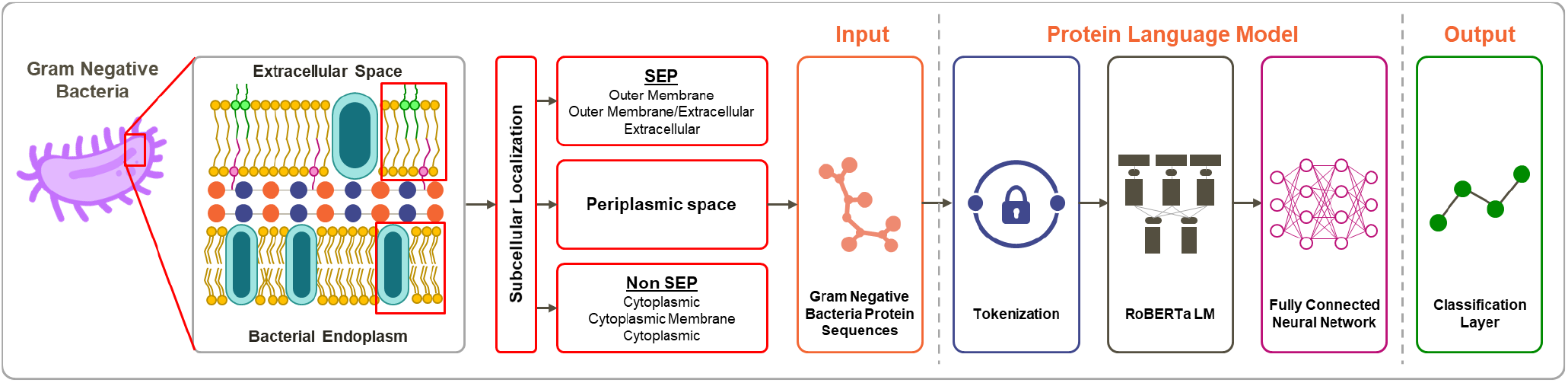
Overview of architecture for ProtLM.SCL: We present a protein language (ProtLM) modelling-based approach using a large dataset of protein amino acid sequences obtained from UniRef50. For the downstream task of predicting subcellular localization (SCL), we appended a fully connected dense layer along with a classification layer to the pretrained general protein language model. This model (ProtLM.SCL) was finetuned and cross-validated on the dataset obtained from Yu et. al. (2010) for two separate tasks: Binary and Multi-class classification.

### Baseline Models

We compared our models to more traditional n-gram-based machine learning approaches to demonstrate the advantage of protein language modelling approach. To accomplish this, we represented bacterial protein sequences as unigram, bigram & trigrams of amino acids strung together with 1 stride. This generated a total of 3 different variations for modelling protein sequences: (i) unigram only (ii) unigram + bigram (iii) unigram + bigram + trigram.

Following that, three different types of models were trained: Logistic Regression (LR), Extreme Gradient Boosting (XGBoost), and Light Gradient Boosting Machines (LightGBM).

## 3. Results

### 3.1. Training and validation of ProtLM.SCL for Binary and Multi-class subcellular localization

For Binary-class classification, ProtLM.SCL achieved an MCC score of 0.92±0.03. It was interesting to see that performance of our model was consistent across all different buckets hence the low score of standard deviation. Upon testing our model on hold-out dataset, the model achieved an MCC score of 0.96±0.01 (**Table S6**) thus assuring that our model was generalizable on unseen data.

For Multi-class classification, we trained individual versions of models each class using one vs. rest methodology. Doing this we achieved class wise MCC scores 0.94±0.02, 0.89±0.02, 0.72±0.08, 0.88±0.05, and 0.74±0.06 for cytoplasmic, cytoplasmic membrane, extracellular, outer membrane and periplasmic respectively (**Table S7-S9**).

Further testing our model on hold-out dataset, the model achieved class wise MCC scores 0.95±0.01, 0.94±0.01, 0.77±0.08, 0.85±0.04, 0.81±0.06 for cytoplasmic, cytoplasmic membrane, extracellular, outer membrane and periplasmic respectively, here also our models were generalized on unseen data (**Table S10-12)**

### 3.2. Baseline performance of ProtLM.SCL for predicting SEP vs. Non-SEP

For establishing baseline, we compared the performance of ProtLM.SCL with three other ML models: LR, LightGBM, and XGboost with n-gram approach considering three different variations (for more details see **section 2.4**). We observed that XGBoost and LightGBM classifier outperformed LR models for cross-validated dataset and their variations achieving an MCC score of 0.83±0.04, 0.83±0.05 and 0.79±0.05 respectively. However, this performance was still inferior to averaged performance metrics for ProtLM.SCL by 11%, 11% and 17% for XGBoost, LightGM and LR respectively, See **Table 1** and **Table S13**.

**Table 1:** Baseline performance of ProtLM.SCL for predicting SEP vs. Non-SEP on cross-validation dataset. We compare the 10-fold cross validation performance of ProtLM.SCL with three alternative baseline N-gram machine learning models: Logistic Regression (LR), Light Gradient Boosting Machines (LightGBM), and Extreme Gradient Boosting Machines (EGBM) (XGBoost). ProtLM.SCL performs better than all n-gram based models by at least 11% - 17% MCC score.

| N GRAM | MODEL | MCC |
| --- | --- | --- |
| Unigram | LR | 0.79±0.05 |
|  | LightGBM | 0.83±0.06 |
|  | XGBoost | 0.82±0.06 |
| Unigram + Bigram | LR | 0.79±0.05 |
|  | LightGBM | 0.83±0.05 |
|  | XGBoost | 0.83±0.04 |
| Unigram + Bigram + Trigram | LR | 0.79±0.05 |
|  | LightGBM | 0.83±0.06 |
|  | XGBoost | 0.82±0.06 |
| Language Model | ProtLM.SCL (10 bucket) | **0.92±0.03** |

Similarly, on the hold-out dataset, ProtLM.SCL outperformed XGBoost, LightGBM and LR Unigram models. The MCC score of XGBoost, LightGBM and LR 0.92±0.01, 0.92±0.01, and 0.88±0.0 respectively. The performance of unigram was better as compared with bigram and trigram model. (**Table S6**).

We have also observed ProtLM.SCL outperformed XGBoost, LightGBM and LR unigram, bigram and trigram models on benchmark dataset (**Table 2**).

**Table 2:** Comparison of the performance of ProtLM.SCL for performing binary classification with other N-gram based ML models on the benchmarking dataset. Our approach outperforms all the three machine learning models and their variants by a wide margin in the comparison (22% - 44% gain of MCC score).

| N GRAM | MODEL | MCC | F1 | ACC | ROC AUC | PRECISION | RECALL |
| --- | --- | --- | --- | --- | --- | --- | --- |
| Unigram | LR | 0.55 | 0.58 | 0.94 | 0.90 | 0.65 | 0.52 |
|  | LightGBM | 0.62 | 0.66 | 0.94 | 0.92 | 0.64 | 0.67 |
|  | XGBoost | 0.59 | 0.63 | 0.93 | 0.91 | 0.62 | 0.64 |
| Unigram + Bigram | LR | 0.56 | 0.59 | 0.94 | 0.91 | 0.66 | 0.53 |
|  | LightGBM | 0.64 | 0.67 | 0.95 | 0.92 | 0.71 | 0.63 |
|  | XGBoost | 0.65 | 0.67 | 0.95 | 0.92 | 0.72 | 0.63 |
| Unigram + Bigram + Trigram | LR | 0.57 | 0.60 | 0.94 | 0.91 | 0.68 | 0.54 |
|  | LightGBM | 0.65 | 0.68 | 0.95 | 0.92 | 0.72 | 0.64 |
|  | XGBoost | 0.64 | 0.67 | 0.95 | 0.92 | 0.73 | 0.61 |
| Language model | ProtLM.SCL | **0.79** | **0.97** | **0.97** | **0.96** | **0.97** | **0.97** |

In conclusion, we observed that our approach outperformed all the three machine learning models by a wide margin. We attribute this improvement in performance to the model’s pretraining on a large dataset of proteins, which may have aided the model in learning complex motifs that are driving protein surface exposition in gram-negative bacteria.

### 3.3. Performance comparison of ProtLM.SCL and PSORTb in predicting SEP vs. non-SEP using benchmarking dataset

We aimed to contrast the performance of PSORTb with ProtLM.SCL on external experimental validated datasets. As detailed in **Table S4** and **Section 2.5**, this was derived from seven experimentally validated datasets^16,21–26^. We were mindful to filter and eliminate the proteins that overlapped between training dataset for our model. This version of the dataset is referred to as the “benchmark dataset” and it contained 3950 proteins (SEP: 339 and non-SEP: 3611). We utilized MCC score to compare performance because it is a more balanced assessment of a classifier^27^.

Here, we have performed five types of comparisons: (i) taking into account all proteins in the benchmarking dataset and treating PSORTb’s ‘Unknown’ predictions as errors since they represent a set of inputs for which the user cannot derive any conclusion; (ii) removing protein sequences predicted as ‘Unknown’ by PSORTb and comparing the performance of ProtLM.SCL on the remaining protein sequences; (iii) removing protein sequences labelled as periplasmic and comparing the performance of ProtLM.SCL on the remaining sequences (iv) removing protein sequences labelled as either ‘Unknown’ and ‘periplasmic’ and comparing the performance of ProtLM.SCL on the remaining sequences (v) Using ProtLM.SCL to predict on sequences predicted as ‘Unknown’ by PSORTb to assess the coverage for our approach. **Figure 4** summarizes these findings.

**Figure 4:**
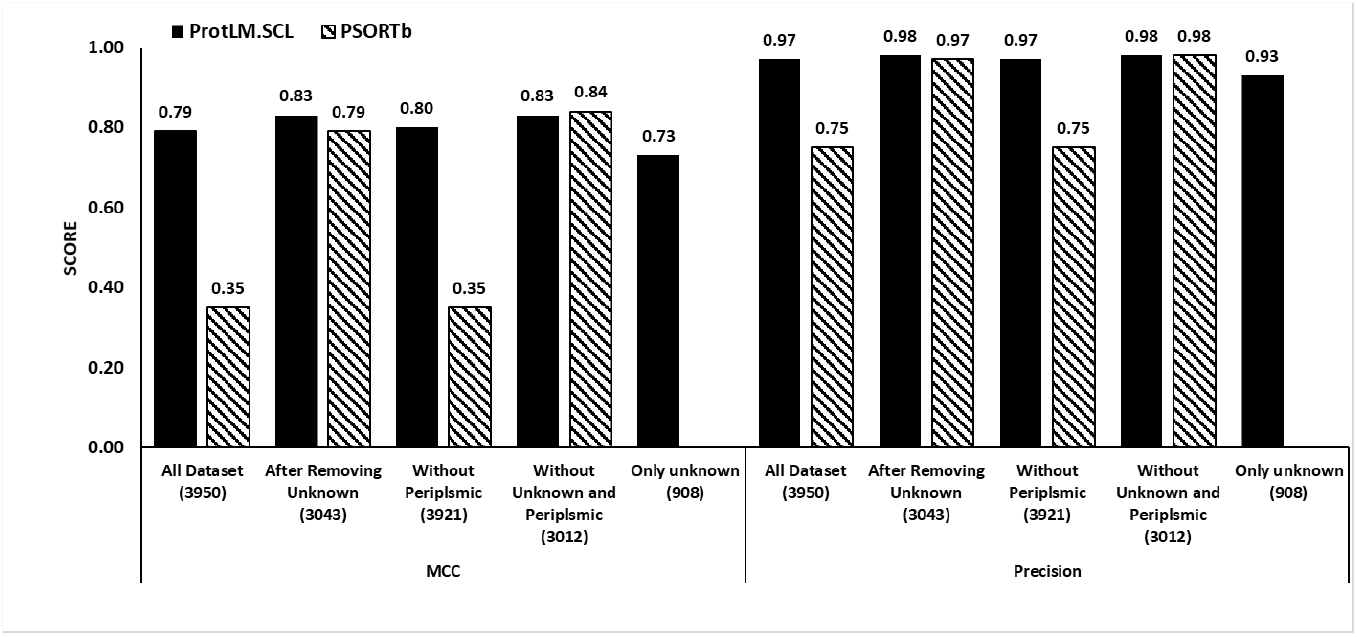
Performance comparison of ProtLM.SCL and PSORTb in predicting SEP vs. non-SEP using benchmarking dataset. We combined the datasets obtained from Sueki et al., 2020, Whitby et al., 2015, Yu et al., 2010, Zhang et al., 2021, Shen and Chou, 2010, Luo, 2012 and Goldberg et al., 2014 to create benchmarking dataset. Thereafter, we have performed five types of comparisons: (i) **All Dataset:** taking into account all proteins in the benchmarking dataset and treating PSORTb’s ‘Unknown’ predictions as errors since they represent a set of inputs for which the user cannot derive any conclusion; (ii) **After Removing ‘Unknown’:** removing protein sequences predicted as ‘Unknown’ by PSORTb and comparing the performance of ProtLM.SCL on the remaining protein sequences; (iii) **Without Periplasmic:** removing protein sequences labelled as periplasmic and comparing the performance of ProtLM.SCL on the remaining sequences (iv) **Without ‘Unknown’ and Periplasmic:** removing protein sequences labelled as either ‘Unknown’ and ‘periplasmic’ and comparing the performance of ProtLM.SCL on the remaining sequences (v) **Only ‘Unknown’**: Using ProtLM.SCL to predict on sequences predicted as ‘Unknown’ by PSORTb to assess the coverage for our approach. The numbers on the X axis indicate number of sequences considered for each type of comparison.

In the first comparison, ProtLM.SCL outperformed PSORTb by 125.7% (ProtLM.SCL: 0.79; PSORTb: 0.35). This is a substantial improvement due to ProtLM.SCL being a more balanced classifier. PSORTb predicted 908 proteins (23%) as unknown in the benchmark dataset.

For the second comparison, wherein we have removed proteins labelled as ‘Unknown’ by PSORTb, the dataset contained a total of 3043 sequences (77% of total sequences). ProtLM.SCL outperformed PSORTb by 5% (ProtLM.SCL: 0.83; PSORTb: 0.79). It was also interesting to note that the performance of ProtLM.SCL also increased by 5% from 0.79 to 0.83 (**Figure 4**).

For the third comparison, upon removing periplasmic sequences, the dataset contained a total of 3921 sequences (99.2% of total sequences). Here, the performance of ProtLM.SCL was significantly better than PSORTb achieving an increase in performance of 129% (ProtLM.SCL 0.80 vs PSORTb 0.35) (**Figure 4**).

For the fourth comparison, upon removing sequences labelled as ‘Unknown’ as predicted by PSORTb or ‘periplasmic’, the dataset contained a total of 3012 sequences (76.2% of total sequences). Here, it was interesting to see that the performance of ProtLM.SCL and PSORTb was comparable (ProtLM.SCL 0.83 vs PSORTb 0.84) (**Figure 4**). It is also important to note here that a significant number of proteins (847 proteins; 93% of total unknown proteins) were predicted correctly by ProtLM.SCL, however not included in this comparison.

Lastly, to test the coverage of ProtLM.SCL, we utilized 908 proteins (23% of total proteins) predicted as ‘Unknown’ by PSORTb for the benchmark dataset. Here, ProtLM.SCL achieved a MCC score 0.73. In conclusion, this offers a major improvement over PSORTb in terms of both performance and coverage (**Figure 4**).

### 3.4. Baseline performance of ProtLM.SCL for predicting multi-class subcellular localization

For multi-class subcellular localization, we performed 10-fold cross validation using ProtLM.SCL to predict the subcellular localization (SCL) on a total of 7,498 proteins obtained from Yu et. al. ^16^, See **Section 2.4** and **Table S3** for more details.

To quantify the baseline performance for SCL categories, we evaluated the performance of LR, XGBoost and LightGBM against ProtLM.SCL. For processing the protein sequence as an input to the baseline models, we created three n-gram-stride versions as described in section 2.6.

We observed similar performances for baseline models across all n-gram variations with XGBoost and LightGBM outperforming LR. ProtLM.SCL performed better than all baseline models across all n-gram variations and in all the categories of subcellular localizations. We observed a best baseline performance of 0.84±0.02 (XGBoost; unigram + bigram + trigram), 0.79±0.03 (LightGBM; unigram + bigram + trigram), 0.62±0.09 (LightGBM; unigram only), 0.68±0.09 (XGBoost; unigram + bigram), and 0.46±0.14 (XGBoost; unigram only), but when we used ProtLM.SCL we observed a performance improvement of 12% (0.94±0.02), 12.6% (0.89±0.02), 16% (0.72±0.08), 30.8% (0.89±0.05) and 60.8% (0.74±0.06) for cytoplasmic, cytoplasmic membrane, extracellular, outer membrane and Periplasmic respectively (see **Table S7-S9**).

Similarly, on the hold-out dataset, ProtLM.SCL outperformed XGBoost, LightGBM and LR by achieving, 0.95±0.01 vs 0.85±0.01 (LightGBM; unigram + bigram + trigram), 0.94±0.01 vs 0.82±0.02 (XGBoost; unigram + bigram + trigram), 0.77±0.08 vs 0.65±0.03 (XGBoost; unigram only), 0.85±0.04 vs 0.83±0.03 (XGBoost; unigram only) and 0.81±0.06 vs 0.57±0.05(LightGBM; unigram only) for cytoplasmic, cytoplasmic membrane, extracellular, outer membrane and Periplasmic respectively (**Table S10-S12**).

We have further checked the performance of ProtLM.SCL along with XGBoost, LightGBM and LR (unigram, unigram + bigram, unigram + bigram + trigram) models on benchmark dataset for each category of subcellular location. ProtLM.SCL performance was better than other n-gram ML models across all the parameters (**Table 3**).

**Table 3:** Comparison of the class-wise performance of ProtLM.SCL for performing multi-class classification with other N-gram based ML models on the benchmarking dataset.

| CATEGORIES | N- GRAM | ML MODEL | MCC | F1 | ACC | ROC_AUC | PRECISION | RECALL |
| --- | --- | --- | --- | --- | --- | --- | --- | --- |
| Cytoplasmic | Unigram | LR | 0.55 | 0.86 | 0.81 | 0.87 | 0.91 | 0.82 |
|  |  | LightGBM | 0.59 | 0.87 | 0.81 | 0.90 | 0.94 | 0.8 |
|  |  | XGBoost | 0.58 | 0.86 | 0.81 | 0.90 | 0.94 | 0.8 |
|  | Unigram + Bigram | LR | 0.54 | 0.86 | 0.8 | 0.88 | 0.92 | 0.81 |
|  |  | LightGBM | 0.60 | 0.87 | 0.82 | 0.91 | 0.94 | 0.81 |
|  |  | XGBoost | 0.59 | 0.86 | 0.81 | 0.91 | 0.94 | 0.80 |
|  | Unigram+ Bigram+ Trigram | LR | 0.54 | 0.86 | 0.80 | 0.88 | 0.92 | 0.80 |
|  |  | LightGBM | 0.61 | 0.87 | 0.83 | 0.91 | 0.95 | 0.81 |
|  |  | XGBoost | 0.61 | 0.87 | 0.82 | 0.92 | 0.95 | 0.81 |
|  | Language Model | ProtLM.SCL | **0.85** | **0.94** | **0.94** | **0.98** | **0.94** | **0.94** |
| Cytoplasmic Membrane | Unigram | LR | 0.66 | 0.69 | 0.94 | 0.88 | 0.74 | 0.65 |
|  |  | LightGBM | 0.61 | 0.65 | 0.92 | 0.91 | 0.59 | 0.72 |
|  |  | XGBoost | 0.59 | 0.63 | 0.92 | 0.91 | 0.58 | 0.70 |
|  | Unigram + Bigram | LR | 0.66 | 0.69 | 0.94 | 0.89 | 0.73 | 0.66 |
|  |  | LightGBM | 0.66 | 0.70 | 0.94 | 0.91 | 0.70 | 0.69 |
|  |  | XGBoost | 0.64 | 0.68 | 0.93 | 0.91 | 0.67 | 0.68 |
|  | Unigram+ Bigram+ Trigram | LR | 0.66 | 0.69 | 0.94 | 0.89 | 0.74 | 0.65 |
|  |  | LightGBM | 0.68 | 0.71 | 0.94 | 0.91 | 0.73 | 0.69 |
|  |  | XGBoost | 0.66 | 0.69 | 0.94 | 0.91 | 0.69 | 0.69 |
|  | Language Model | ProtLM.SCL | **0.89** | **0.98** | **0.98** | **0.98** | **0.98** | **0.98** |
| Extracellular | Unigram | LR | 0.27 | 0.29 | 0.96 | 0.82 | 0.34 | 0.25 |
|  |  | LightGBM | 0.45 | 0.47 | 0.97 | 0.85 | 0.51 | 0.43 |
|  |  | XGBoost | 0.50 | 0.52 | 0.97 | 0.85 | 0.55 | 0.49 |
|  | Unigram +<br>Bigram | LR | 0.28 | 0.3 | 0.96 | 0.82 | 0.35 | 0.26 |
|  |  | LightGBM | 0.50 | 0.5 | 0.97 | 0.88 | 0.62 | 0.42 |
|  |  | XGBoost | 0.48 | 0.48 | 0.97 | 0.87 | 0.58 | 0.42 |
|  | Unigram+<br>Bigram+ Trigram | LR | 0.28 | 0.3 | 0.96 | 0.83 | 0.35 | 0.26 |
|  |  | LightGBM | 0.46 | 0.46 | 0.97 | 0.88 | 0.59 | 0.38 |
|  |  | XGBoost | 0.52 | 0.52 | 0.97 | 0.87 | 0.62 | 0.45 |
|  | Language Model | ProtLM.SCL | **0.57** | **0.97** | **0.97** | **0.91** | **0.97** | **0.97** |
| Outer<br>Membrane | Unigram | LR | 0.20 | 0.21 | 0.95 | 0.88 | 0.32 | 0.16 |
|  |  | LightGBM | 0.46 | 0.47 | 0.96 | 0.89 | 0.59 | 0.39 |
|  |  | XGBoost | 0.45 | 0.46 | 0.96 | 0.88 | 0.57 | 0.39 |
|  | Unigram +<br>Bigram | LR | 0.20 | 0.21 | 0.95 | 0.88 | 0.33 | 0.15 |
|  |  | LightGBM | 0.49 | 0.47 | 0.97 | 0.90 | 0.73 | 0.34 |
|  |  | XGBoost | 0.52 | 0.49 | 0.97 | 0.91 | 0.79 | 0.36 |
|  | Unigram+<br>Bigram+ Trigram | LR | 0.21 | 0.21 | 0.95 | 0.88 | 0.34 | 0.15 |
|  |  | LightGBM | 0.52 | 0.50 | 0.97 | 0.90 | 0.79 | 0.36 |
|  |  | XGBoost | 0.45 | 0.43 | 0.97 | 0.91 | 0.69 | 0.31 |
|  | Language Model | ProtLM.SCL | **0.81** | **0.98** | **0.98** | **0.99** | **0.98** | **0.98** |
| Periplasmic | Unigram | LR | 0.15 | 0.11 | 0.92 | 0.84 | 0.47 | 0.06 |
|  |  | LightGBM | 0.36 | 0.36 | 0.93 | 0.83 | 0.59 | 0.26 |
|  |  | XGBoost | 0.38 | 0.35 | 0.93 | 0.81 | 0.68 | 0.24 |
|  | Unigram +<br>Bigram | LR | 0.17 | 0.12 | 0.93 | 0.85 | 0.51 | 0.07 |
|  |  | LightGBM | 0.39 | 0.35 | 0.94 | 0.85 | 0.77 | 0.22 |
|  |  | XGBoost | 0.36 | 0.32 | 0.93 | 0.84 | 0.72 | 0.21 |
|  | Unigram+<br>Bigram+ Trigram | LR | 0.17 | 0.12 | 0.93 | 0.85 | 0.53 | 0.07 |
|  |  | LightGBM | 0.33 | 0.26 | 0.93 | 0.86 | 0.78 | 0.16 |
|  |  | XGBoost | 0.38 | 0.33 | 0.94 | 0.85 | 0.78 | 0.21 |
|  | Language Model | ProtLM.SCL | **0.70** | **0.96** | **0.96** | **0.91** | **0.96** | **0.96** |

### 3.5. Performance comparison of ProtLM.SCL and PSORTb for predicting multi-class localization using benchmarking datasets

We performed three kinds of comparisons: (i) Considering all proteins from all data sets and treating ‘Unknown’ predictions from PSORTb and ProtLM.SCL’s as errors; (ii) removing the protein set predicted as ‘Unknown’ by PSORTb/ProtLM.SCL and comparing their predictions; (iii) To assess the coverage of both models, we considered the sequences predicted as ‘Unknown’ by either models and performed a comparative analysis using PSORTb or ProtLM.SCL.

For the first comparison, the combined performance (MCC and precision) of ProtLM.SCL outperformed PSORTb. The percentage gain of ProtLM.SCL in MCC and precision was 60.07% and 32.51% respectively (**Figure 5**). Furthermore, we have evaluated and compared the class wise MCC and precision scores of ProtLM.SCL and PSORTb. We observed the difference in performance for MCC scores of +63.50%, +85.89%, +129.09%, +150.24%, +107.20% and precision score of +32.35%, +33%, +28.80%, +28.49%, and +27.79% when comparing ProtLM.SCL with PSORTb for cytoplasmic, cytoplasmic membrane, periplasmic, outer membrane and extracellular categories respectively (**Table S14**).

**Figure 5:**
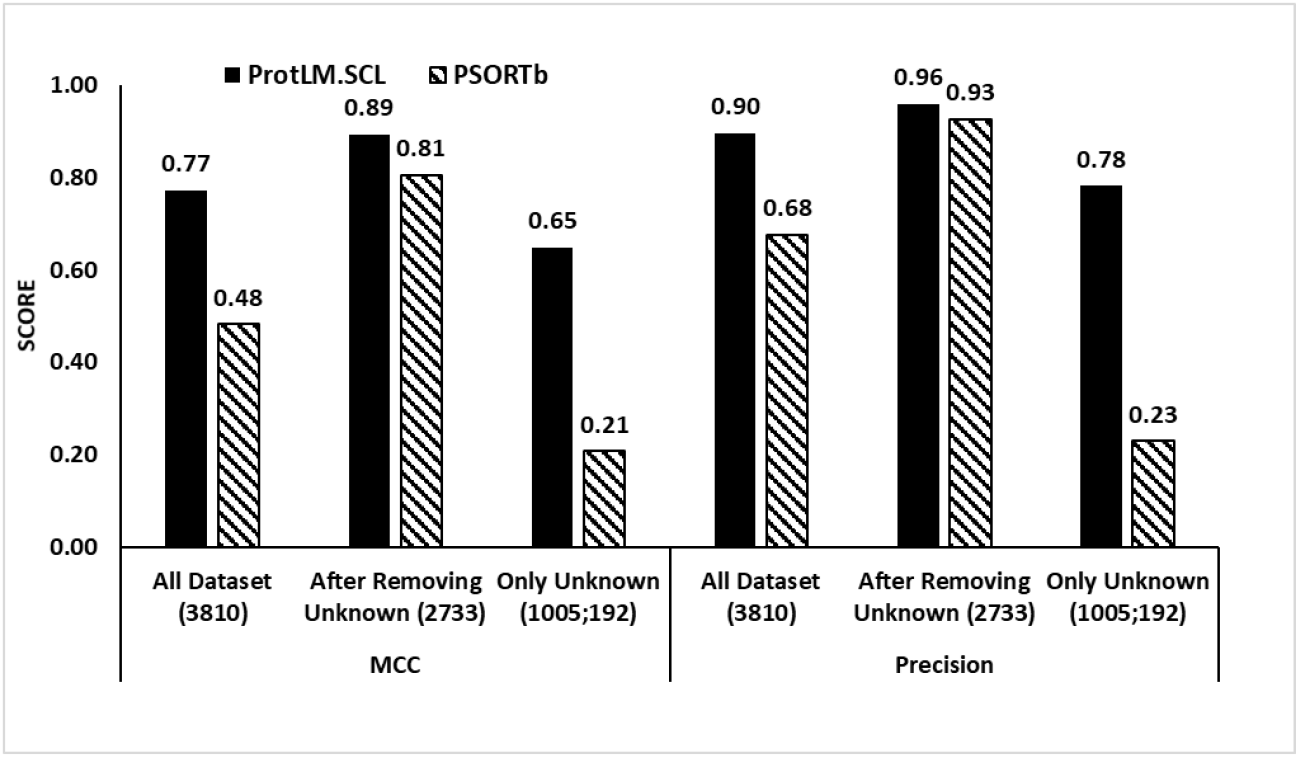
Performance comparison of ProtLM.SCL and PSORTb for predicting multi-class localization using benchmarking datasets. We report the combined performance for ProtLM.SCL and PSORTb on 5 subcellular localizations i.e. Cytoplasm, Cytoplasmic Membrane, Extracellular, Outer Membrane, Periplasmic. To accomplish multi-class classification, we trained five distinct instances of the language model, one for each of the five target labels using one vs. rest strategy, namely Cytoplasmic, Cytoplasmic Membrane, Periplasmic, Outer Membrane, and Extracellular. To determine the final localization, we use the highest probability score derived from the output of the five models. Additionally, if we none of the models achieved a probability of >0.5, the prediction for such protein an input is considered an error and labelled as ‘Unknown’. The numbers in X axis indicate number of sequences considered in benchmark dataset. We performed three kinds of comparisons: (i) Considering all proteins from all data sets and treating PSORTb and ProtLM.SCL’s ‘Unknown’ predictions as errors; (ii) removing the protein set predicted as ‘Unknown’ by PSORTb/ProtLM.SCL and comparing their predictions; (iii) To assess the coverage of both models, we considered the sequences predicted as ‘Unknown’ by both PSORTb and ProtLM.SCL. The numbers on the X axis indicate number of sequences considered in each type of comparison.

For the second comparison, we removed the proteins labelled as ‘Unknown’ by PSORTb and ProtLM.SCL, we were left with 2733 protein sequences. The overall percentage gain by ProtLM.SCL for MCC and precision was 10.8% and 3.75% respectively (**Figure 5**). Here, we observed the percentage gain in performance for class wise MCC +8.65%, +14.42%, +13.49%, +4.65%, +2.80% and precision +2.82%, +2.83%, +0.68%, +0.11%, and 0.0% when comparing ProtLM.SCL with PSORTb for cytoplasmic, cytoplasmic membrane, periplasmic, outer membrane and extracellular categories respectively (**Table S14**).

For assessing the coverage of models, a total of 1005 protein sequences (26.3% of total proteins) were labelled as ‘Unknown’ by PSORTb while 192 sequences (5.03% of total proteins) were labelled as ‘Unknown’ by ProtLM.SCL. Out of 192 sequences, 120 sequences were overlapping with the ‘Unknown’ predicted by PSORTb. When testing ProtLM.SCL on protein sequences predicted as ‘Unknown’ by PSORTb, we observed an averaged MCC score of 0.65 and precision score of 0.78 for ProtLM.SCL (**Figure 5**). The class wise performance of ProtLM.SCL on this category is summarized in **Table S14**. In constrast, PSORTb performed poorly in the ‘Unknown’ ProtLM.SCL category (**Figure 5 and Table S14**)

## Discussion

Subcellular localization of gram-negative bacterial proteins can be determined using a variety of experimental approaches such as immunolabeling, fractionation, and others. However, these processes can be laborious, time consuming and costly, particularly in the early phases of drug discovery, when the pool of candidates is often relatively large^28^. As a result, computational prediction models have gained prominence in the post-genomic era, for determining the subcellular localization and molecular interactions. This includes tools to predict the global localization of bacterial proteins, such as PSORTb, which predicts the SCL of bacterial proteins, and the Signal P server, which uses predictive algorithms to spot the presence and location of signal peptides or cleavage sites in prokaryotic and eukaryotic peptide sequences^29,30^. However, the usability of these tools is hindered by performance limitation and/or due to the large number of proteins labelled as ‘Unknown’ i.e. missing label or prediction.

To address these limitations, we present a Protein Language Modeling-based approach for predicting the subcellular localization (ProtLM.SCL) of gram-negative bacteria proteins, see **Figure 1**. This model was trained and validated using publicly available experimental data^16,21–26^. We generated two types of ProtLM.SCL models for gram-negative bacterial proteins: (i) Binary classification, in which we classified outer membrane and extracellular as potential SEPs, while cytoplasmic and cytoplasmic membrane were classified as non-SEPs. (ii) Multi-class classification for predicting five subcellular localizations: cytoplasmic, cytoplasmic membrane, periplasmic, outer membrane, and extracellular. We trained these models on the PSORTb dataset and then benchmarked their predictive performance using several comparisons: (i) comparison with other baseline machine learning models such as LR, LightGBM, and XGBoost (ii) Testing the performances on PSORTb’s ‘Unknown’ predictions, which represent a set of inputs for which the user cannot derive any conclusion (iii) comparison with PSORTb.

To summarize, we present a unique language modeling-based approach for predicting subcellular localization of gram-negative bacterial proteins. We determined that our strategy had a number of advantages (i) **Simplified input:** our model requires only the primary sequences of proteins as an input which makes it widely useful for any protein localization prediction task (ii) **Improved performance:** When compared to baseline models and PSORTb, we consistently observed a gain of performance in MCC score for binary and multi-class classifications, (iii) **Greater coverage:** When predicting for ‘Unknown’ labels from PSORTb, we discovered that ProtLM.SCL recovered considerable amount of these predictions.

One limitation of this version is that of our binary classification model was not trained on the periplasmic proteins. By manually annotating this subset of proteins into SEP and non-SEP labels, we expect an increase of performances. Additionally, addressing the interpretability of these models to highlight key regions responsible for the model predictions. This model is specific for gram-negative bacteria and cannot be extended *per se* to other types.

## Conclusion

In conclusion, we demonstrated that by using protein language modelling, we can train models that accurately predict gram-negative bacterial protein subcellular localization (ProtLM.SCL). Our model outperformed not just n-gram-based machine learning models, but also the most widely used tools such as PSORTb in terms of performance and coverage. Furthermore, in addition to developing a multi-class classification model, we present a particular case of developing a binary model that is tuned toward direct prediction of protein surface exposure, which is the desired outcome in reverse vaccinology applications for antigen discovery. These models are likely to be critical in the creation of vaccine targets and the investigation of bacterial-host interactions, and in this regard, ProtLM.SCL represents a substantial advancement over currently employed methodologies.

## Sponsorship and Acknowledgements

This work was sponsored by GSK which was involved in all stages of the study conduct and analysis. We thank Alex Pysik, Andy Bonner, Guglielmo Roma, Janet Effler, Kalyani Anantapantula, Matt Harrison, Mara Peccanti, Rebecca Stephens, Sai Jasti, Paul Smyth, Alexander(Dan) Hudson, Nathalie Norais, Alessandro Brozzi and Yannick Vanloubbeeck for their support and valuable inputs.

