## Supplementary file for "Protein Language Model for Prediction of Subcellular Localization of Protein Sequences from Gram-negative bacteria (ProtLM.SCL)"

### Supplementary Information

#### S1.Supplementary Tables

| S.No. | Subcellular localization | Count | Binary class | Multi-class |
|---|---|---|---|---|
| 1 | Cytoplasmic | 5025 | Non-SEP | Cytoplasmic |
| 2 | Cytoplasmic/Cytoplasmic Membrane | 50 | Non-SEP | Cytoplasmic and Cytoplasmic Membrane |
| 3 | Cytoplasmic Membrane | 1607 | Non-SEP | Cytoplasmic Membrane |
| 4 | Cytoplasmic Membrane/Periplasmic | 58 | | Cytoplasmic Membrane and Periplasmic |
| 5 | Periplasmic | 437 | | Periplasmic |
| 6 | Periplasmic/Outer Membrane | 3 | | Periplasmic and Outer Membrane |
| 7 | Outer Membrane | 541 | SEP | Outer Membrane |
| 8 | Outer Membrane/Extracellular | 90 | SEP | Outer Membrane and Extracellular |
| 9 | Extracellular | 419 | SEP | Extracellular |

**Table S1:** Distribution of the protein subcellular localization for the cross-validation dataset.

| Data Split | Bucket | PROTEIN COUNTS | SEP % |
|---|---|---|---|
| Cross Validation | 1 | 665 | 13 |
| | 2 | 673 | 14 |
| | 3 | 687 | 17 |
| | 4 | 682 | 17 |
| | 5 | 701 | 13 |
| | 6 | 704 | 12 |
| | 7 | 718 | 14 |
| | 8 | 711 | 11 |
| | 9 | 747 | 14 |
| | 10 | 747 | 11 |
| **Hold-Out** | 11 | 691 | 14 |

**Table S2:** Protein count and distribution of SCL in the buckets for cross-validation dataset for binary classification.

| Data Split | Bucket | #RECORDS | % CYTOPLASMIC | % CYTOPLASMIC MEMBRANE | % PERIPLASMIC | % OUTER MEMBRANE | % EXTRACELLULAR |
|---|---|---|---|---|---|---|---|
| Cross-Validation | 1 | 724 | 63 | 18 | 8 | 8 | 6 |
| | 2 | 723 | 63 | 18 | 7 | 7 | 7 |
| | 3 | 730 | 58 | 20 | 6 | 10 | 8 |
| | 4 | 739 | 55 | 22 | 8 | 10 | 7 |
| | 5 | 745 | 65 | 19 | 6 | 7 | 7 |
| | 6 | 751 | 62 | 22 | 6 | 7 | 5 |
| | 7 | 765 | 63 | 20 | 6 | 8 | 5 |
| | 8 | 752 | 63 | 22 | 5 | 6 | 5 |
| | 9 | 788 | 61 | 23 | 5 | 8 | 6 |
| | 10 | 781 | 67 | 19 | 4 | 6 | 5 |
| **Hold-out** | 11 | 726 | 58 | 25 | 5 | 7 | 7 |

**Table S3:** Protein count and distribution of SEP in the buckets for cross-validation dataset for Multiclass classification.

| S.NO. | DATASET SOURCE | SEP/NON-SEP | SCL | # ORGANISM PER DATASET |
|---|---|---|---|---|
| **1** | Sueki et al., 2020 | 2767 | 2614 | 1 |
| **2** | Whitby et al., 2015 | 53 | NA | 1 |
| **3** | Yu et al., 2010 | 162 | 162 | 1 |
| **4** | Zhang et al., 2021 | 61 | 64 | 1 |
| **5** | Shen and Chou, 2010 | 51 | 53 | 42 |
| **6** | Luo, 2012 | 88 | 96 | 46 |
| **7** | Goldberg et al., 2014 | 768 | 821 | 526 |
| | **Total** | **3950** | **3810** | |

**Table S4:** Information about distribution of proteins to perform binary and/or multi-class classification on benchmarking dataset compiled from Sueki et al., 2020, Whitby et al., 2015, Yu et al., 2010, Zhang et al., 2021, Shen and Chou, 2010, Luo, 2012 and Goldberg et al., 2014 along with organism counts for each dataset.

| **Subcellular localization from publications and Swiss-Prot** | **ProtLM.SCL renamed Subcellular Localization** |
|---|---|
| Secreted, cell wall, S-layer | Extracellular |
| ‘Cytoplasm’<br>‘Cytoplasm, cytosol’<br>'Cytoplasmic vesicle' | Cytoplasmic |
| Periplasm<br>'Periplasmic flagellum', | Periplasmic |
| ‘Cell membrane’,<br>‘Cell inner membrane; Single-pass membrane protein; Periplasmic side’,<br>‘Cell inner membrane; Multi-pass membrane protein; Periplasmic side’,<br>‘Cell inner membrane; Multi-pass membrane protein’,<br>‘Cell inner membrane,<br>Cell inner membrane; Multi-pass membrane protein; Cytoplasmic side’,<br>'Cell inner membrane; Single-pass type II membrane protein',<br>'Cell inner membrane; Single-pass type I membrane protein',<br>'Cell inner membrane; Single-pass type I membrane protein; Periplasmic side' | Cytoplasmic Membrane |
| 'Cell outer membrane; Single-pass membrane protein',<br>'Cell outer membrane ',<br>'Cell outer membrane; Multi-pass membrane protein' | Outer Membrane |

**Table S5: Mapping of subcellular localization Labels.** Basic data cleansing was performed, and subcellular categories were standardized as per PSORTb nomenclature.

| N GRAM | MODEL | MCC | PRECISION | ACC | F1 | RECALL | ROC_AUC |
|---|---|---|---|---|---|---|---|
| Unigram | LR | 0.88±0.00 | 0.99±0.00 | 0.97±0.00 | 0.89±0.00 | 0.80±0.01 | 0.99±0.00 |
| | LightGBM | 0.92±0.01 | 0.99±0.01 | 0.98±0.00 | 0.93±0.01 | 0.88±0.01 | 1.00±0.00 |
| | XGBoost | 0.92±0.01 | 0.99±0.01 | 0.98±0.00 | 0.93±0.01 | 0.88±0.02 | 1.00±0.00 |
| Unigram + Bigram | LR | 0.87±0.00 | 0.99±0.00 | 0.97±0.00 | 0.89±0.00 | 0.80±0.00 | 0.99±0.00 |
| | LightGBM | 0.91±0.01 | 1.00±0.00 | 0.98±0.00 | 0.92±0.01 | 0.85±0.01 | 1.00±0.00 |
| | XGBoost | 0.91±0.01 | 1.00±0.00 | 0.98±0.00 | 0.92±0.01 | 0.85±0.02 | 1.00±0.00 |
| Unigram + Bigram + Trigram | LR | 0.87±0.00 | 0.99±0.00 | 0.97±0.00 | 0.89±0.00 | 0.80±0.00 | 0.99±0.00 |
| | LightGBM | 0.91±0.01 | 1.00±0.00 | 0.98±0.00 | 0.91±0.01 | 0.84±0.01 | 1.00±0.00 |
| | XGBoost | 0.92±0.02 | 1.00±0.01 | 0.98±0.00 | 0.92±0.01 | 0.86±0.02 | 1.00±0.00 |
| **Language Model** | ProtLM.SCL | **0.96±0.01** | **0.99±0.00** | **0.99±0.00** | **0.99±0.00** | **0.99±0.00** | **1.00±0.00** |

**Table S6**: **Comparison of performance between ProtLM.SCL and simpler ML models for predicting SEP vs. Non-SEP on cross-validation on Hold-out set.**

| LOCALIZATION | MODEL | ACC | F1 | MCC | PRECISION | RECALL | ROC_AUC |
|---|---|---|---|---|---|---|---|
| **Cytoplasmic** | LR | 0.88±0.02 | 0.91±0.01 | 0.74±0.04 | 0.88±0.03 | 0.94±0.01 | 0.94±0.02 |
| | LightGBM | 0.92±0.01 | 0.94±0.01 | 0.83±0.03 | 0.92±0.02 | 0.95±0.02 | 0.97±0.01 |
| | XGBoost | 0.92±0.01 | 0.93±0.01 | 0.82±0.03 | 0.92±0.02 | 0.95±0.02 | 0.97±0.01 |
| | PROTLM.SCL | **0.97±0.01** | **0.97±0.01** | **0.94±0.02** | **0.97±0.01** | **0.98±0.01** | **0.99±0.00** |
| **Cytoplasmic Membrane** | LR | 0.92±0.01 | 0.77±0.05 | 0.75±0.04 | 0.96±0.02 | 0.64±0.07 | 0.91±0.02 |
| | LightGBM | 0.93±0.01 | 0.81±0.03 | 0.77±0.04 | 0.92±0.04 | 0.72±0.05 | 0.94±0.01 |
| | XGBoost | 0.93±0.01 | 0.80±0.04 | 0.77±0.04 | 0.92±0.04 | 0.72±0.05 | 0.94±0.01 |
| | PROTLM.SCL | **0.96±0.00** | **0.91±0.01** | **0.89±0.02** | **0.95±0.03** | **0.87±0.03** | **0.98±0.01** |
| **Extracellular** | LR | 0.95±0.00 | 0.47±0.08 | 0.50±0.06 | 0.82±0.09 | 0.34±0.07 | 0.94±0.02 |
| | LightGBM | 0.96±0.01 | 0.62±0.10 | 0.62±0.09 | 0.80±0.10 | 0.51±0.10 | 0.96±0.01 |
| | XGBoost | 0.96±0.00 | 0.62±0.10 | 0.62±0.10 | 0.79±0.11 | 0.51±0.11 | 0.96±0.01 |
| | PROTLM.SCL | **0.97±0.01** | **0.72±0.08** | **0.72±0.08** | **0.82±0.15** | **0.67±0.13** | **0.97±0.02** |
| **Outer Membrane** | LR | 0.93±0.02 | 0.41±0.09 | 0.41±0.09 | 0.64±0.12 | 0.30±0.09 | 0.95±0.02 |
| | LightGBM | 0.96±0.01 | 0.69±0.08 | 0.68±0.08 | 0.82±0.09 | 0.60±0.10 | 0.96±0.01 |
| | XGBoost | 0.96±0.01 | 0.67±0.08 | 0.66±0.08 | 0.81±0.09 | 0.59±0.10 | 0.96±0.01 |
| | PROTLM.SCL | **0.98±0.01** | **0.89±0.05** | **0.88±0.05** | **0.89±0.07** | **0.89±0.06** | **0.99±0.01** |
| **Periplasmic** | LR | 0.94±0.01 | 0.15±0.09 | 0.22±0.11 | 0.67±0.25 | 0.09±0.06 | 0.88±0.04 |
| | LightGBM | 0.95±0.01 | 0.44±0.10 | 0.45±0.11 | 0.70±0.14 | 0.32±0.09 | 0.91±0.04 |
| | XGBoost | 0.95±0.01 | 0.44±0.14 | 0.46±0.14 | 0.70±0.16 | 0.33±0.13 | 0.92±0.03 |
| | PROTLM.SCL | **0.97±0.01** | **0.75±0.06** | **0.74±0.06** | **0.78±0.09** | **0.73±0.06** | **0.97±0.01** |

**Table S7: A comparison of the performance between ProtLM.SCL and LR, XGBoost, and LightGBM with protein sequences transposed as unigram input for achieving multi-class classification using 10-fold cross validation strategy.** Protein sequences were split into unigram and fed into LR, LightGBM, and XGBoost as inputs, see section 2.6 for more details.

| LOCALIZATION | MODEL | ACC | F1 | MCC | PRECISION | RECALL | ROC_AUC |
|---|---|---|---|---|---|---|---|
| **Cytoplasmic** | LR | 0.89±0.02 | 0.91±0.01 | 0.76±0.03 | 0.89±0.02 | 0.94±0.01 | 0.95±0.01 |
| | LightGBM | 0.92±0.01 | 0.94±0.01 | 0.83±0.02 | 0.92±0.02 | 0.95±0.01 | 0.97±0.01 |
| | XGBoost | 0.92±0.01 | 0.94±0.01 | 0.83±0.02 | 0.92±0.02 | 0.96±0.01 | 0.97±0.01 |
| | PROTLM.SCL | **0.97±0.01** | **0.97±0.01** | **0.94±0.02** | **0.97±0.01** | **0.98±0.01** | **0.99±0.00** |
| **Cytoplasmic Membrane** | LR | 0.93±0.01 | 0.78±0.04 | 0.76±0.04 | 0.97±0.01 | 0.66±0.06 | 0.92±0.01 |
| | LightGBM | 0.93±0.01 | 0.81±0.03 | 0.79±0.03 | 0.96±0.02 | 0.71±0.06 | 0.95±0.01 |
| | XGBoost | 0.93±0.01 | 0.82±0.03 | 0.79±0.03 | 0.96±0.02 | 0.71±0.06 | 0.95±0.01 |
| | PROTLM.SCL | **0.96±0.00** | **0.91±0.01** | **0.89±0.02** | **0.95±0.03** | **0.87±0.03** | **0.98±0.01** |
| **Extracellular** | LR | 0.96±0.00 | 0.47±0.08 | 0.51±0.06 | 0.82±0.09 | 0.34±0.07 | 0.95±0.02 |
| | LightGBM | 0.96±0.01 | 0.53±0.11 | 0.55±0.11 | 0.82±0.16 | 0.40±0.09 | 0.96±0.01 |
| | XGBoost | 0.96±0.01 | 0.55±0.10 | 0.57±0.10 | 0.83±0.14 | 0.42±0.10 | 0.96±0.01 |
| | PROTLM.SCL | **0.97±0.01** | **0.72±0.08** | **0.72±0.08** | **0.82±0.15** | **0.67±0.13** | **0.97±0.02** |
| **Outer Membrane** | LR | 0.93±0.02 | 0.44±0.09 | 0.44±0.09 | 0.68±0.12 | 0.33±0.09 | 0.95±0.02 |
| | LightGBM | 0.96±0.01 | 0.67±0.10 | 0.67±0.09 | 0.86±0.07 | 0.56±0.13 | 0.97±0.01 |
| | XGBoost | 0.96±0.01 | 0.67±0.10 | 0.68±0.09 | 0.87±0.08 | 0.56±0.12 | 0.97±0.01 |
| | PROTLM.SCL | **0.98±0.01** | **0.89±0.05** | **0.88±0.05** | **0.89±0.07** | **0.89±0.06** | **0.99±0.01** |
| **Periplasmic** | LR | 0.94±0.01 | 0.16±0.09 | 0.23±0.11 | 0.68±0.26 | 0.09±0.06 | 0.89±0.04 |
| | LightGBM | 0.95±0.01 | 0.32±0.14 | 0.38±0.13 | 0.78±0.18 | 0.21±0.11 | 0.92±0.04 |
| | XGBoost | 0.95±0.01 | 0.35±0.10 | 0.41±0.09 | 0.82±0.07 | 0.22±0.07 | 0.92±0.04 |
| | PROTLM.SCL | **0.97±0.01** | **0.75±0.06** | **0.74±0.06** | **0.78±0.09** | **0.73±0.06** | **0.97±0.01** |

**Table S8: A comparison of the performance between ProtLM.SCL and LR, XGBoost, and LightGBM with protein sequences transposed as unigram + bigram input for achieving multi-class classification using 10-fold cross validation strategy.** Protein sequences were split into unigram, bigram and fed together into LR, LightGBM, and XGBoost as inputs, see section 2.6 for more details.

| LOCALIZATION | MODEL | ACC | F1 | MCC | PRECISION | RECALL | ROC_AUC |
|---|---|---|---|---|---|---|---|
| **Cytoplasmic** | LR | 0.89±0.02 | 0.92±0.01 | 0.77±0.03 | 0.89±0.02 | 0.94±0.01 | 0.95±0.01 |
| | LightGBM | 0.92±0.01 | 0.94±0.01 | 0.84±0.02 | 0.92±0.02 | 0.96±0.01 | 0.98±0.01 |
| | XGBoost | 0.92±0.01 | 0.94±0.01 | 0.84±0.02 | 0.93±0.02 | 0.95±0.01 | 0.98±0.01 |
| | PROTLM.SCL | **0.97±0.01** | **0.97±0.01** | **0.94±0.02** | **0.97±0.01** | **0.98±0.01** | **0.99±0.00** |
| **Cytoplasmic Membrane** | LR | 0.93±0.01 | 0.79±0.04 | 0.77±0.04 | 0.97±0.01 | 0.66±0.06 | 0.92±0.01 |
| | LightGBM | 0.94±0.01 | 0.82±0.04 | 0.79±0.03 | 0.97±0.02 | 0.71±0.06 | 0.96±0.01 |
| | XGBoost | 0.94±0.01 | 0.82±0.03 | 0.79±0.03 | 0.96±0.02 | 0.71±0.05 | 0.95±0.01 |
| | PROTLM.SCL | **0.96±0.00** | **0.91±0.01** | **0.89±0.02** | **0.95±0.03** | **0.87±0.03** | **0.98±0.01** |
| **Extracellular** | LR | 0.96±0.00 | 0.47±0.07 | 0.51±0.06 | 0.83±0.09 | 0.34±0.07 | 0.95±0.02 |
| | LightGBM | 0.96±0.01 | 0.53±0.10 | 0.55±0.11 | 0.82±0.15 | 0.39±0.08 | 0.96±0.02 |
| | XGBoost | 0.96±0.01 | 0.55±0.08 | 0.57±0.08 | 0.86±0.13 | 0.41±0.07 | 0.96±0.01 |
| | PROTLM.SCL | **0.97±0.01** | **0.72±0.08** | **0.72±0.08** | **0.82±0.15** | **0.67±0.13** | **0.97±0.02** |
| **Outer Membrane** | LR | 0.94±0.02 | 0.44±0.08 | 0.45±0.08 | 0.69±0.12 | 0.34±0.09 | 0.95±0.02 |
| | LightGBM | 0.96±0.01 | 0.66±0.11 | 0.66±0.09 | 0.88±0.07 | 0.54±0.13 | 0.97±0.01 |
| | XGBoost | 0.96±0.01 | 0.66±0.10 | 0.67±0.09 | 0.87±0.07 | 0.55±0.13 | 0.97±0.01 |
| | PROTLM.SCL | **0.98±0.01** | **0.89±0.05** | **0.88±0.05** | **0.89±0.07** | **0.89±0.06** | **0.99±0.01** |
| **Periplasmic** | LR | 0.94±0.01 | 0.15±0.10 | 0.22±0.13 | 0.63±0.33 | 0.09±0.06 | 0.89±0.04 |
| | LightGBM | 0.95±0.01 | 0.29±0.12 | 0.35±0.12 | 0.79±0.19 | 0.18±0.08 | 0.93±0.03 |
| | XGBoost | 0.95±0.01 | 0.34±0.10 | 0.41±0.09 | 0.82±0.11 | 0.22±0.08 | 0.92±0.04 |
| | PROTLM.SCL | **0.97±0.01** | **0.75±0.06** | **0.74±0.06** | **0.78±0.09** | **0.73±0.06** | **0.97±0.01** |

**Table S9: A comparison of the performance between ProtLM.SCL and LR, XGBoost, and LightGBM with protein sequences transposed as unigram + bigram + trigram input for achieving multi-class classification using 10-fold cross validation strategy.** Protein sequences were split into unigram, bigram and trigram and fed together into LR, LightGBM, and XGBoost as inputs, see section 2.6 for more details.

| LOCALIZATION | MODEL | MCC | PRECISION | ACC | F1 | RECALL | ROC_AUC |
|---|---|---|---|---|---|---|---|
| **Cytoplasmic** | LR | 0.74±0.00 | 0.86±0.00 | 0.87±0.00 | 0.90±0.00 | 0.94±0.00 | 0.93±0.00 |
| | LightGBM | 0.83±0.01 | 0.91±0.01 | 0.92±0.01 | 0.93±0.00 | 0.95±0.00 | 0.97±0.00 |
| | XGBoost | 0.82±0.01 | 0.90±0.00 | 0.91±0.00 | 0.93±0.00 | 0.95±0.00 | 0.97±0.00 |
| | ProtLM.SCL | **0.95±0.01** | **0.98±0.0** | **0.98±0.0** | **0.98±0.0** | **0.98±0.0** | **0.99±0.00** |
| **CytoplasmicMembrane** | LR | 0.75±0.00 | 0.98±0.00 | 0.91±0.00 | 0.78±0.00 | 0.65±0.00 | 0.90±0.00 |
| | LightGBM | 0.79±0.01 | 0.95±0.01 | 0.92±0.00 | 0.83±0.01 | 0.73±0.01 | 0.96±0.00 |
| | XGBoost | 0.78±0.02 | 0.94±0.02 | 0.92±0.01 | 0.82±0.01 | 0.73±0.01 | 0.96±0.00 |
| | ProtLM.SCL | **0.94±0.01** | **0.98±0.0** | **0.98±0.0** | **0.98±0.01** | **0.98±0.0** | **0.98±0.0** |
| **Extracellular** | LR | 0.40±0.01 | 0.63±0.01 | 0.94±0.00 | 0.40±0.01 | 0.30±0.01 | 0.97±0.00 |
| | LightGBM | 0.64±0.02 | 0.74±0.03 | 0.96±0.00 | 0.66±0.02 | 0.59±0.03 | 0.97±0.00 |
| | XGBoost | 0.65±0.03 | 0.75±0.04 | 0.96±0.00 | 0.67±0.03 | 0.60±0.03 | 0.97±0.00 |
| | ProtLM.SCL | **0.77±0.08** | **0.97±0.01** | **0.97±0.01** | **0.97±0.01** | **0.97±0.01** | **0.99±0.01** |
| **OuterMembrane** | LR | 0.55±0.02 | 0.73±0.03 | 0.95±0.00 | 0.56±0.01 | 0.45±0.01 | 0.96±0.00 |
| | LightGBM | 0.81±0.03 | 0.91±0.03 | 0.98±0.00 | 0.82±0.03 | 0.75±0.03 | 0.99±0.00 |
| | XGBoost | 0.83±0.03 | 0.92±0.04 | 0.98±0.00 | 0.84±0.03 | 0.77±0.04 | 0.99±0.00 |
| | ProtLM.SCL | **0.85±0.04** | **0.98±0.01** | **0.98±0.01** | **0.98±0.01** | **0.98±0.01** | **0.99±0.00** |
| **Periplasmic** | LR | 0.22±0.03 | 0.61±0.05 | 0.95±0.00 | 0.16±0.03 | 0.09±0.02 | 0.89±0.00 |
| | LightGBM | 0.57±0.05 | 0.83±0.08 | 0.97±0.00 | 0.54±0.05 | 0.41±0.05 | 0.95±0.01 |
| | XGBoost | 0.55±0.07 | 0.82±0.11 | 0.97±0.00 | 0.53±0.06 | 0.39±0.05 | 0.95±0.01 |
| | ProtLM.SCL | **0.81±0.06** | **0.98±0.01** | **0.98±0.01** | **0.98±0.01** | **0.98±0.01** | **0.99±0.01** |

**Table S10: A comparison of the performance between ProtLM.SCL, and LR, XGBoost, and LightGBM with protein sequences transposed as unigram input for achieving multi-class classification using 10-fold cross validation on Hold-out set.** Protein sequences were split into unigram and fed into LR, LightGBM, and XGBoost as inputs, see section 2.6 for more details.

| LOCALIZATION | MODEL | MCC | PRECISION | ACC | F1 | RECALL | ROC_AUC |
|---|---|---|---|---|---|---|---|
| **Cytoplasmic** | LR | 0.76±0.01 | 0.87±0.00 | 0.88±0.00 | 0.90±0.00 | 0.94±0.00 | 0.94±0.00 |
| | LightGBM | 0.83±0.01 | 0.91±0.01 | 0.92±0.01 | 0.93±0.00 | 0.95±0.01 | 0.97±0.00 |
| | XGBoost | 0.83±0.02 | 0.91±0.01 | 0.92±0.01 | 0.93±0.01 | 0.95±0.01 | 0.97±0.00 |
| | **ProtLM.SCL** | **0.95±0.01** | **0.98±0.0** | **0.98±0.0** | **0.98±0.0** | **0.98±0.0** | **0.99±0.00** |
| **CytoplasmicMembrane** | LR | 0.75±0.00 | 0.97±0.00 | 0.91±0.00 | 0.78±0.00 | 0.65±0.00 | 0.92±0.00 |
| | LightGBM | 0.81±0.01 | 0.97±0.01 | 0.93±0.00 | 0.84±0.01 | 0.75±0.01 | 0.97±0.00 |
| | XGBoost | 0.80±0.01 | 0.97±0.01 | 0.93±0.00 | 0.84±0.01 | 0.74±0.02 | 0.96±0.00 |
| | **ProtLM.SCL** | **0.94±0.01** | **0.98±0.0** | **0.98±0.0** | **0.98±0.01** | **0.98±0.0** | **0.98±0.00** |
| **Extracellular** | LR | 0.41±0.02 | 0.61±0.01 | 0.94±0.00 | 0.41±0.02 | 0.31±0.02 | 0.97±0.00 |
| | LightGBM | 0.57±0.04 | 0.70±0.03 | 0.95±0.00 | 0.59±0.04 | 0.51±0.05 | 0.97±0.00 |
| | XGBoost | 0.57±0.05 | 0.69±0.06 | 0.95±0.01 | 0.59±0.05 | 0.52±0.06 | 0.96±0.00 |
| | **ProtLM.SCL** | **0.77±0.08** | **0.97±0.01** | **0.97±0.01** | **0.97±0.01** | **0.97±0.01** | **0.99±0.01** |
| **OuterMembrane** | LR | 0.57±0.01 | 0.73±0.01 | 0.95±0.00 | 0.58±0.01 | 0.48±0.01 | 0.96±0.00 |
| | LightGBM | 0.80±0.03 | 0.91±0.01 | 0.97±0.00 | 0.81±0.03 | 0.73±0.05 | 0.98±0.00 |
| | XGBoost | 0.80±0.03 | 0.90±0.04 | 0.97±0.00 | 0.81±0.02 | 0.73±0.02 | 0.98±0.00 |
| | **ProtLM.SCL** | **0.85±0.04** | **0.98±0.01** | **0.98±0.01** | **0.98±0.01** | **0.98±0.01** | **0.99±0.00** |
| **Periplasmic** | LR | 0.27±0.01 | 0.68±0.04 | 0.95±0.00 | 0.20±0.00 | 0.11±0.00 | 0.89±0.00 |
| | LightGBM | 0.53±0.03 | 0.94±0.05 | 0.97±0.00 | 0.47±0.03 | 0.31±0.02 | 0.95±0.01 |
| | XGBoost | 0.55±0.04 | 0.94±0.07 | 0.97±0.00 | 0.49±0.05 | 0.33±0.05 | 0.95±0.01 |
| | **ProtLM.SCL** | **0.81±0.06** | **0.98±0.01** | **0.98±0.01** | **0.98±0.01** | **0.98±0.01** | **0.99±0.01** |

**Table S11: A comparison of the performance between ProtLM.SCL, and LR, XGBoost, and LightGBM with protein sequences transposed as unigram + bigram input for achieving multi-class classification using 10-fold cross validation on Hold-out set.** Protein sequences were split into unigram and bigram and fed together into LR, LightGBM, and XGBoost as inputs, see section 2.6 for more details.

| LOCALIZATION | MODEL | MCC | PRECISION | ACC | F1 | RECALL | ROC_AUC |
|---|---|---|---|---|---|---|---|
| **Cytoplasmic** | LR | 0.77±0.00 | 0.87±0.00 | 0.89±0.00 | 0.91±0.00 | 0.95±0.00 | 0.94±0.00 |
| | LightGBM | 0.85±0.01 | 0.92±0.00 | 0.93±0.00 | 0.94±0.00 | 0.96±0.00 | 0.98±0.00 |
| | XGBoost | 0.85±0.01 | 0.92±0.00 | 0.93±0.01 | 0.94±0.00 | 0.95±0.01 | 0.97±0.00 |
| | **ProtLM.SCL** | **0.95±0.01** | **0.98±0.0** | **0.98±0.0** | **0.98±0.0** | **0.98±0.0** | **0.99±0.00** |
| **CytoplasmicMembrane** | LR | 0.75±0.00 | 0.97±0.01 | 0.91±0.00 | 0.78±0.00 | 0.65±0.00 | 0.92±0.00 |
| | LightGBM | 0.81±0.01 | 0.99±0.01 | 0.93±0.00 | 0.83±0.01 | 0.72±0.02 | 0.97±0.00 |
| | XGBoost | 0.82±0.02 | 0.98±0.01 | 0.93±0.01 | 0.85±0.02 | 0.75±0.02 | 0.97±0.00 |
| | **ProtLM.SCL** | **0.94±0.01** | **0.98±0.0** | **0.98±0.0** | **0.98±0.01** | **0.98±0.0** | **0.98±0.0** |
| **Extracellular** | LR | 0.43±0.03 | 0.63±0.02 | 0.94±0.00 | 0.43±0.03 | 0.33±0.03 | 0.97±0.00 |
| | LightGBM | 0.55±0.03 | 0.69±0.02 | 0.95±0.00 | 0.56±0.03 | 0.48±0.03 | 0.97±0.00 |
| | XGBoost | 0.55±0.07 | 0.67±0.06 | 0.95±0.01 | 0.57±0.07 | 0.51±0.07 | 0.96±0.01 |
| | **ProtLM.SCL** | **0.77±0.08** | **0.97±0.01** | **0.97±0.01** | **0.97±0.01** | **0.97±0.01** | **0.99±0.01** |
| **OuterMembrane** | LR | 0.57±0.02 | 0.72±0.01 | 0.95±0.00 | 0.58±0.02 | 0.49±0.02 | 0.96±0.00 |
| | LightGBM | 0.77±0.02 | 0.90±0.02 | 0.97±0.00 | 0.78±0.02 | 0.69±0.03 | 0.98±0.00 |
| | XGBoost | 0.79±0.01 | 0.91±0.02 | 0.97±0.00 | 0.79±0.01 | 0.70±0.01 | 0.98±0.00 |
| | **ProtLM.SCL** | **0.85±0.04** | **0.98±0.01** | **0.98±0.01** | **0.98±0.01** | **0.98±0.01** | **0.99±0.00** |
| **Periplasmic** | LR | 0.27±0.01 | 0.69±0.06 | 0.95±0.00 | 0.20±0.00 | 0.11±0.00 | 0.89±0.00 |
| | LightGBM | 0.49±0.05 | 0.87±0.04 | 0.96±0.00 | 0.43±0.06 | 0.29±0.05 | 0.93±0.01 |
| | XGBoost | 0.50±0.05 | 0.86±0.07 | 0.96±0.00 | 0.45±0.05 | 0.31±0.04 | 0.95±0.01 |
| | **ProtLM.SCL** | **0.81±0.06** | **0.98±0.01** | **0.98±0.01** | **0.98±0.01** | **0.98±0.01** | **0.99±0.01** |

**Table S12: A comparison of the performance between ProtLM.SCL, and LR, XGBoost, and LightGBM with protein sequences transposed as unigram + bigram + trigram input for achieving multi-class classification using 10-fold cross validation on Hold-out set.** Protein sequences were split into unigram, bigram and trigram and fed together into LR, LightGBM, and XGBoost as inputs, see section 2.6 for more details.

| N GRAM | MODEL | ACC | F1 | PRECISION | RECALL | ROC_AUC |
|---|---|---|---|---|---|---|
| **Unigram** | **LR** | 0.95±0.01 | 0.80±0.05 | 0.94±0.04 | 0.70±0.07 | 0.97±0.01 |
| | **LightGBM** | 0.96±0.01 | 0.85±0.05 | 0.91±0.06 | 0.81±0.06 | 0.98±0.01 |
| | **XGBoost** | 0.96±0.01 | 0.84±0.05 | 0.91±0.06 | 0.78±0.07 | 0.98±0.01 |
| **Unigram + Bigram** | **LR** | 0.95±0.01 | 0.81±0.05 | 0.94±0.03 | 0.71±0.07 | 0.98±0.01 |
| | **LightGBM** | 0.96±0.01 | 0.85±0.05 | 0.94±0.03 | 0.77±0.07 | 0.98±0.01 |
| | **XGBoost** | 0.96±0.01 | 0.84±0.04 | 0.94±0.03 | 0.77±0.06 | 0.98±0.01 |
| **Unigram + Bigram + Trigram** | **LR** | 0.95±0.01 | 0.80±0.05 | 0.94±0.04 | 0.70±0.07 | 0.97±0.01 |
| | **LightGBM** | 0.96±0.01 | 0.85±0.05 | 0.91±0.06 | 0.81±0.06 | 0.98±0.01 |
| | **XGBoost** | 0.96±0.01 | 0.84±0.05 | 0.91±0.06 | 0.78±0.07 | 0.98±0.01 |
| **Language Model** | **ProtLM.SCL (10bucket)** | **0.98±0.01** | **0.98±0.00** | **0.98 ±0.01** | **0.98 ±0.01** | **0.96 ±0.01** |

**Table S13**: **Baseline performance of ProtLM.SCL for predicting SEP vs. Non-SEP on cross-validation dataset.** We compare the 10-fold cross validation performance of ProtLM.SCL with three alternative baseline N-gram machine learning models: Logistic Regression (LR), Light Gradient Boosting Machines (LightGBM), and Extreme Gradient Boosting Machines (EGBM) (XGBoost).

| DATASETS | SUBCELLULAR CATEGORIES | PROTLM.SCL | | | | | PSORTB | | | | |
|---|---|---|---|---|---|---|---|---|---|---|---|
| | | MCC | PRECISION | F1 | RECALL | ACC | MCC | PRECISION | F1 | RECALL | ACC |
| ALL DATASET | **Cytoplasmic** | 0.8044 | 0.9181 | 0.9417 | 0.9181 | 0.9181 | 0.492 | 0.6937 | 0.7984 | 0.6937 | 0.6937 |
| | **Cytoplasmic Membrane** | 0.7274 | 0.931 | 0.9553 | 0.931 | 0.931 | 0.3913 | 0.7000 | 0.8076 | 0.7000 | 0.7000 |
| | **Periplasmic** | 0.5663 | 0.9236 | 0.9457 | 0.9236 | 0.9236 | 0.2472 | 0.7171 | 0.8194 | 0.7171 | 0.7171 |
| | **Outer Membrane** | 0.5788 | 0.9362 | 0.9611 | 0.9362 | 0.9362 | 0.2313 | 0.7286 | 0.8379 | 0.7286 | 0.7286 |
| | **Extracellular** | 0.4142 | 0.9207 | 0.9467 | 0.9207 | 0.9207 | 0.1999 | 0.7205 | 0.83 | 0.7205 | 0.7205 |
| AFTER REMOVING UNKNOWN | **Cytoplasmic** | 0.9248 | 0.9744 | 0.9743 | 0.9744 | 0.9744 | 0.8512 | 0.9477 | 0.9482 | 0.9477 | 0.9477 |
| | **Cytoplasmic Membrane** | 0.9165 | 0.9824 | 0.9826 | 0.9824 | 0.9824 | 0.801 | 0.9554 | 0.9569 | 0.9554 | 0.9554 |
| | **Periplasmic** | 0.786 | 0.9828 | 0.9826 | 0.9828 | 0.9828 | 0.6926 | 0.9762 | 0.9754 | 0.9762 | 0.9762 |
| | **Outer Membrane** | 0.8647 | 0.9916 | 0.9919 | 0.9916 | 0.9916 | 0.8263 | 0.9905 | 0.9904 | 0.9905 | 0.9905 |
| | **Extracellular** | 0.7229 | 0.9817 | 0.9822 | 0.9817 | 0.9817 | 0.7032 | 0.9817 | 0.9816 | 0.9817 | 0.9817 |
| PSORTb's Unknown | **Cytoplasmic** | 0.6924 | 0.8308 | 0.8841 | 0.8308 | 0.8308 | - | - | - | - | - |
| | **Cytoplasmic Membrane** | 0.5025 | 0.8577 | 0.913 | 0.8577 | 0.8577 | - | - | - | - | - |
| | **Periplasmic** | 0.5044 | 0.8289 | 0.8774 | 0.8289 | 0.8289 | - | - | - | - | - |
| | **Outer Membrane** | 0.5115 | 0.8527 | 0.9086 | 0.8527 | 0.8527 | - | - | - | - | - |
| | **Extracellular** | 0.2316 | 0.8209 | 0.8844 | 0.8209 | 0.8209 | - | - | - | - | - |
| ProtLM.SCL Unknown | **Cytoplasmic** | - | - | - | - | - | 0.1853 | 0.276 | 0.3966 | 0.276 | 0.276 |
| | **Cytoplasmic Membrane** | - | - | - | - | - | 0.1084 | 0.2917 | 0.4343 | 0.2917 | 0.2917 |
| | **Periplasmic** | - | - | - | - | - | 0.1533 | 0.3333 | 0.472 | 0.3333 | 0.3333 |
| | **Outer Membrane** | - | - | - | - | - | 0.1242 | 0.3594 | 0.5233 | 0.3594 | 0.3594 |
| | **Extracellular** | - | - | - | - | - | 0.1107 | 0.3229 | 0.4684 | 0.3229 | 0.3229 |

**Table S14**: **Summary of the multi-class classification with details regarding category wise performance on benchmark dataset using ProtLM.SCL and PSORTb from class-wise models.** Please note that due to unavailability of probability score for predictions from PSORTb, ROC-AUC could not be calculated, hence the metric has been excluded for category wise performance matrix.

#### S2. Detailed information regarding the benchmarking datasets

##### *S2.1. Dataset obtained from Goldberg et al., 2014*

We downloaded the LocTREE3 bacterial protein dataset (4898 bacterial proteins) which are experimentally annotated of single localization (Swiss-Prot release 2013_11). After removing the gram-positive protein sequences from the dataset, we are left with 2854 gram-negative proteins sequences. Further we checked their subcellular localization of the gram-negative bacterial protein sequences in Swiss-Prot. Based on the subcellular localization names we mapped the localization names to the following category (Table **S 5**).

After preprocessing and removing the overlapping sequences from the training and other benchmark datasets, we considered 768 and 821 protein sequences for SEP/Non-SEP and SCL predictions respectively.

##### *S2.2. Dataset obtained from Shen and Chou, 2010*

Gneg-mPLoc datasets (1392 proteins) were retrieved. After preprocessing and removing the overlapped sequences, we got 51 and 53 sequences for SEP vs Non-SEP and SCL predictions respectively. We mapped the localization names of Gneg-PLoc such as Cell Inner membrane to cytoplasmic membrane, cell outer membrane to outer membrane, cytoplasm to cytoplasmic and extracell to extracellular.

##### *S2.3.* Dataset obtained from Zhang et al., 2021

We retrieved 69 proteins from the publication (Zhang et al., 2021). After pre-processing and removing the sequences which are part of the training dataset, we considered 61 and 64 proteins for SEP/Non-SEP and SCL prediction respectively.

##### *S2.4. Dataset obtained from Luo, 2012*

Out of 1444 proteins, we got 104 proteins which are not part of training dataset. After pre-processing, we considered 88 sequences for SEP/Non-SEP and 96 for SCL prediction.

##### *S2.5. Dataset obtained from Sueki et al., 2020*

We have retrieved 4272 protein sequences from this dataset. After validating their repetition and pre-processing, we concluded with 2767 protein sequences for SEP/Non-SEP and 2614 proteins for SCL predictions

##### *S2.6. Dataset obtained from Whitby et al., 2015*

We considered 53 protein sequences out of 56 sequences from this dataset for SEP/Non-SEP prediction and none of the sequences are considered for SCL prediction since their proper localization labels were not available.

##### *S2.7. Dataset obtained from Yu et al., 2010*

In the PSORTb benchmark dataset, we got 162 proteins which are not part of training dataset. After pre-processing, we considered 162 sequences for Non-SEP as all of them are cytoplasmic and same sequence for SCL prediction.
